# Microbiome network connectivity and composition linked to disease resistance in strawberry plants

**DOI:** 10.1101/2022.10.07.511207

**Authors:** M. Amine Hassani, Omar Gonzalez, Samuel S. Hunter, Gerald J. Holmes, Shashika S. Hewavitharana, Kelly Ivors, Cristina Lazcano

## Abstract

Plant recruit diverse microbial communities from the soil biota. Inter-microbial interactions and connectivity in the root microbiome could play essential roles in plant health by promoting resistance to soil-borne pathogens. Yet, understanding these interactions under field conditions is still scarce. Using a strawberry crop model, we characterized the prokaryotic and fungal communities in the rhizosphere and roots of three strawberry cultivars displaying varying resistance degrees to the soil-borne fungal pathogen *Macrophomina phaseolina.* We tested the hypothesis that resistant cultivars assemble distinct bacterial and fungal communities that foster microbial connectivity and mediate disease resistance. Our results show that the soil-borne pathogen, *M. phaseolina*, does not perturb the root microbiome of the strawberry cultivars. Microbiome comparative analysis indicated that the highly susceptible cultivar, Sweet Ann, assembles a distinct microbiome that shows reduced network connectivity, whereas more resistant cultivars were enriched in potential beneficial microbes and showed higher network connectivity. Collectively, these results suggest the role of plant genetic traits in the assembly of beneficial microbiome members. Our study reinforces the eminent role of the plant microbiome as trait of selection in breeding programs and stresses further understandings of the genetic and biological mechanisms that mediate microbiome assembly. Uncovering these mechanisms will be key for future plant breeding programs.

## Introduction

Plants are colonized by diverse and multi-kingdom microbial communities, collectively designated as the plant microbiome. These microbe associates extend functional repertoires of plants and together with their host form the plant holobiont. Plant-microbiome interactions are at the essence of plant growth and overall health. Harnessing these interactions could be a path forward to expand sustainable agriculture practices. Nevertheless, our understanding of these interactions under field conditions is still limited and requires more empirical studies (Qu et al. 2020; Munoz Ucros et al. 2021).

Strawberries (*Fragaria* × *ananassa* Duchesne ex Rozier) is one of the most recently domesticated garden plants with a little more than 300 years of breeding history (Hardigan et al. 2021). Strawberry fruit are highly appreciated in many areas of the world and often constitute a component of a healthy diet due to their high content of vitamins, phytochemicals and bioactive compounds (Hannum 2004). To control soil-borne fungal pathogens of strawberries, growers over recent decades have relied on the application of fumigants, such as methyl bromide and chloropicrin (Holmes et al. 2020). However, fumigant use has either been eliminated, as in the case of methyl bromide, or increasingly regulated due to their deleterious effects on human and environmental health which coerced growers to search for alternatives, such as breeding new resistant cultivars (Lloyd 2016; Holmes et al. 2020).

The soil-borne pathogen, *Macrophomina phaseolina*, is a plurivorous fungus that infects more than 500 host plants, including the popular garden strawberry (Marquez et al. 2021). This fungus causes dry rot root, also known as charcoal rot root, and remains persistent in infested soils for several years (Short 1980; Su et al. 2001; Gupta et al. 2012). *M. phaseolina* is widely distributed geographically and thrives in high temperature (30-35 °C) and low moisture soils (Marquez et al. 2021). The phaseout of the fumigant methyl bromide coupled with negative effects of global warming (rising temperatures and extended drought periods) renders *M. phaseolina* a global concern as it is expected to reach further geographical regions, expand its natural host range and causes more severe outbreaks (Zveibil et al. 2012; Pickel et al. 2020; Pandey and Basandrai 2021; Cohen et al. 2022). Although there are not known host-resistance genes, breeding plant cultivars that are less susceptible to the pathogen remains a preferred disease management strategy (Holmes et al. 2020; Basandrai et al. 2021; Cohen et al. 2022). In fact, plant cultivars resistant to *M. phaseolina* are tolerant to the infection, as these plants could only limit, and not suppress, the fungal infection or they compensate for the burden of the infection on the plant by increased fitness (Marquez et al. 2021). Plant cultivars with varying degrees of resistance to *M. phaseolina* have been successfully identified in soybean, sesame and strawberry (Reznikov et al. 2018; Gomez et al. 2020; Nelson et al. 2021). Most of our understanding of molecular mechanisms underlying resistance to *M. phaseolina* infection is still focused on identifying defense-related genes and proteins expressed upon infection (Marquez et al. 2021; Kaur et al. 2012). In contrary, a limited number of studies have investigated the role of the plant microbiome in mediating disease resistance against *M. phaseolina* infection (Chowdhury et al. 2014; Lazcano et al. 2021). Recent research has suggested the role of the rhizosphere microbiome of modern strawberry cultivars in mediating resistance against fungal soil-borne pathogens, such as *Verticillium dahliae* and *M. phaseolina* (Lazcano et al. 2021). It is well known that the soil is the main microbial source of rhizosphere and root microbiomes (Bulgarelli et al. 2012; Lundberg et al. 2012). Hence, the capacity of plants to recruit beneficial microbial associates, that some could mediate disease resistance, is tightly related to their genetic traits (Escudero-Martinez et al. 2022; Oyserman et al. 2022). Here, we evaluated the response of the rhizosphere and root microbiomes of three strawberry cultivars (i.e. Sweet Ann, Marquis and Manresa) to soil infestation by the fungal pathogen *M. phaseolina.* The strawberry cultivars, Sweet Ann, Marquis, and Manresa, were shown previously to have high, moderate and low susceptibility to *M. phaseolina*, respectively. We hypothesized that the rhizosphere and the root microbiomes of resistant cultivars assemble distinct bacterial and fungal communities that foster microbial connectivity and mediate disease resistance. To test this, we conducted a comparative microbiome and network analyses and revealed patterns in the assembly of the bacterial and fungal microbiome in the rhizosphere and root microbiome of the three strawberry cultivars. Our study emphasizes the role of rhizosphere and root microbiome as a potential strategy for breeding resistant cultivars to soil-borne pathogens.

## Results

### The root microbiome of strawberry remains unperturbed by the soil-borne pathogen *Macrophomina phaseolina*

We tested the impact of the soil-borne pathogen, *Macrophomina phaseolina*, on the composition and the structure of the root and the rhizosphere microbiota of three strawberry cultivars (i.e. Manresa, Marquis and Sweet Ann). Bare-root transplants from these three cultivars were transplanted into the field and manually inoculated with *M. phaseolina* and grown in parallel to control plants that were not inoculated. Bulk soil, rhizosphere and roots samples were harvested at flowering and harvest, 8 and 14 weeks after transplanting, respectively, for a total of 16 plants per cultivar. To reveal the overall impact of *M. phaseolina* on the rhizosphere and the root microbiomes of strawberries, we have aggregated all time points for each plant cultivar separately. By comparing alpha-diversity measures, we did not observe any significant alteration of the microbial evenness (Bacteria: **Fig 1A**, Fungi: **Fig 1B**) nor richness (Bacteria: **Supplementary Fig 1A**, Fungi: **Supplementary Fig 1B**) due to the presence of the pathogen. However, an exception is noted for the bacterial root microbiome of Manresa. Soil infestation by *M. phaseolina* did negatively, but moderately, alter the evenness and the richness of root bacterial microbiome of Manresa (**Fig 1A and Supplementary Fig 1A, respectively**). These results indicate that the polyphagous pathogen, *M. phaseolina*, has a minor impact on the composition of the rhizosphere and the roots microbial communities of strawberry plants.

**Fig. 1.**
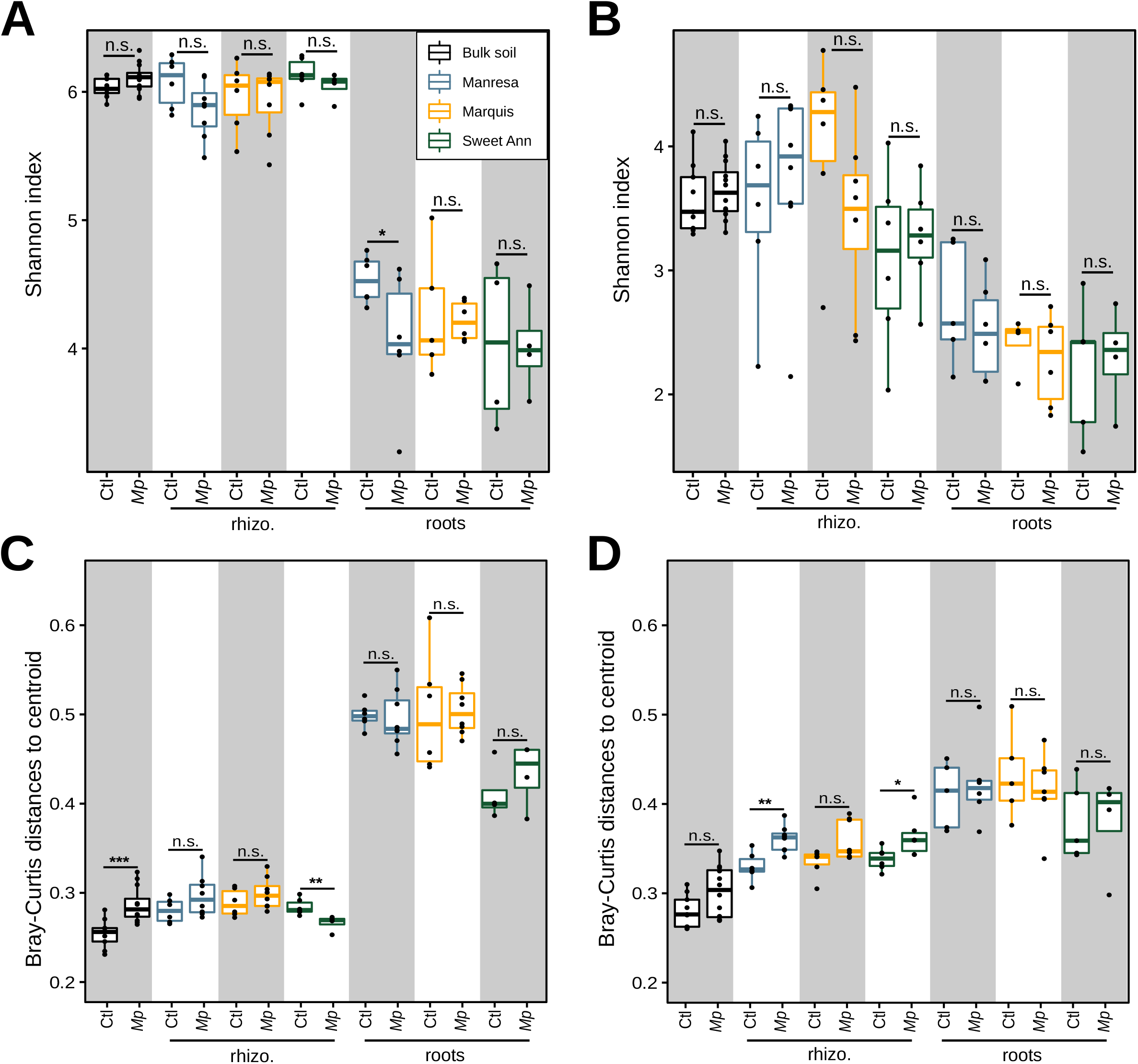
Composition and structure of the bacterial and fungal microbiome of strawberry cultivars upon *Macrophomina phaseolina* infestation. **A** and **B**, show box-plots that depict the alpha diversity measure, Shannon index, of bacterial and fungal communities associated with different strawberry cultivars, respectively. Overall, *M. phaseolina* did not have a significant impact the diversity bacterial and fungal microbiome in the bulk soil, rhizosphere and root of all test strawberry cultivars, except root bacterial microbiome of Manresa that showed slight decrease in diversity. Samples were rarefied to 1000 reads and significance difference was testing using Kruskal-Wallis test. **C** and **D** show BC distance to centroid that indicate community homogeneity. Overall, the community homogeneity was not significantly altered for both bacterial and fungal communities, except increased BC distance of the bacterial communities associated to the bulk soil upon *M. phaseolina* infestation and decreased BC distances in the rhizosphere bacterial microbiome of Sweet Ann. We note a trend of increased BC distance in the rhizosphere fungal microbiome that was significant in Manresa. Color-code is common over the plots and denotes strawberry cultivar.Reads were normalized using cumulative sum scaling prior to computing BC distances and significance difference was testing using Kruskal-Wallis test.

To further test whether this pathogen alters the structure of the microbial communities associated with the rhizosphere and roots of strawberry plants, we performed permutational multivariate analysis of variances (PERMANOVA) (Anderson 2001; Warton et al. 2012) to show enriched bacterial and fungal ASVs that mediate the shifts in the community structure (see **Methods**). These analyses revealed that soil infestation by *M. phaseolina* strongly alters the bacterial and the fungal microbial communities of the bulk soil (Bacteria: **Table 1** “R^2^=0.181, p-value 0.001”**, Supplementary Fig 1C, Supplementary-table 1**, Fungi: **Table 1** “R^2^=0.171, p-value=0.001”**, Supplementary Fig 1D, Supplementary-table 1)** and the rhizosphere fungal microbiome of all test strawberry cultivars (**Table 1** “Manresa, R^2^=0.166, p-value=0.002; Marquis, R^2^=0.135, p-value=0.003 and Sweet Ann R^2^=0.158, p-value=0.004”, **Supplementary Fig 2B**). The observed shifts in the rhizosphere fungal microbiome were mainly mediated by the depletion of several ASVs belonging to the genus *Penicillium*, known to include also plant-growth promoting species (Babu et al. 2015), and the enrichment of a fungal ASV corresponding to *M. phaseolina*. (**Supplementary Fig 2B**). In contrast to bulk soil and the rhizosphere fungal microbiomes, only the rhizosphere bacterial microbiome of Manresa was significantly altered by *M. phaseolina* (**Table 1** “Manresa, R^2^=0.152, p-value=0.009”). In this case, fungal infestation led to the depletion of mainly four bacterial ASVs, two *Burkholderiales* (*Caenimonas sp, Massillia sp)* and two *Solirubrobacterales* (*Conexibacter sp, Solirubrobacter sp*, **Supplementary Fig 2A**). Despite the alteration of the bulk soil and rhizosphere microbial communities, we report only a marginal alteration of the root fungal microbiome and no significant alteration of the root bacterial microbiome (**Table 1** “roots”). Furthermore, the analysis of Bray-Curtis (BC) distances to centroid indicates that *M. phaseolina* did not induce significant heterogeneity in the bacterial and fungal community structures (**Fig 1E** and **1F**, respectively), with exceptions of increased BC distances to centroid of bulk soil bacterial communities (**Fig 1E**) and rhizosphere fungal communities associated with Manresa, both upon *M. phaseolina* infestation (**Fig 1F**). Taken together, our results show that the soil-borne pathogen, *M. phaseolina*, has only a minor impact on the composition and the structure of the bacterial and the fungal root microbiomes of strawberry and further suggest that these microbial communities are rather resistant to this fungal invasion.

**Table1.**
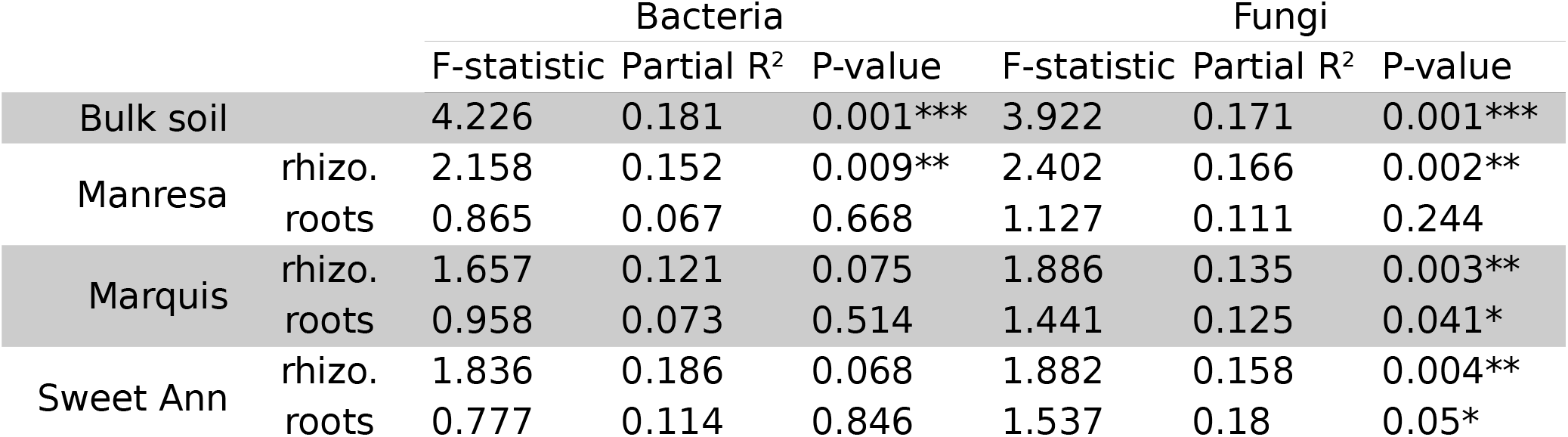
Permutational multivariate analysis of variances in Bray-Curtis distances. Prior to testing, reads were normalized using cumulative sum scaling. Starts indicate significant differences between the control versus M. phaseolina infested samples for corresponding habitat (i.e. bulk soil, rhizosphere or roots).

### Resistant strawberry cultivars assemble a distinct microbiome

The three strawberry cultivars, Manresa, Marquis and Sweet Ann were previously reported to have low, intermediate and high susceptibility to the soil-borne pathogen *M. phaseolina*, respectively (Lazcano et al. 2021). By evaluating percent plant survival at the end of the experiment, we have further corroborated previous findings and showed that Sweet Ann is highly susceptible, Marquis is intermediately susceptible, and Manresa is lowly susceptible (more resistant) to the soil-borne fungal pathogen *M. phaseolina* (**Fig 2A**). Although the two strawberry cultivars Manresa and Marquis showed higher survival rates than Sweet Ann, we detected *M. phaseolina* in the root tissues of all three strawberry cultivars (**Fig 2B**, “Early”, “Late”). These results indicate that *M. phaseolina* is able to infect both cultivars irrespective of their susceptibility rates. Nonetheless, the biomass of *M. phaseolina* was significantly higher at harvest in the cultivar Sweet Ann than in Manresa and Marquis (**Fig 2B** “Late, p-value = 0.03”). The current results prompt us to test whether the three strawberry cultivars assemble distinct rhizosphere and root microbiomes as they showed distinct susceptibility rates to *M. phaseolina*. To this end, we computed BC distances between all samples and displayed these distances using constrained principal coordinate analysis (Anderson and Willis 2003). The constrained analysis was performed by the interaction terms: plant cultivar (i.e. Manresa, Marquis or Sweet Ann), treatment (i.e. infested with *M. phaseolina* or control non-infested) and sample type (i.e. rhizosphere or roots sample). This analysis shows that the assembly of bacterial and fungal communities were strongly driven by sample type (Bacteria: **Fig 2C**, CAP1 29%, F-statistic=32.96, p-value=0.001, triangle *vs* square; Fungi: **Fig 2D**, CAP1 17.1%, F-statistic=16.71, p-value=0.001, triangle *vs* square), as demonstrated in previous plant model systems (Bulgarelli et al. 2012; Lundberg et al. 2012). Notably, the rhizosphere and the root bacterial microbiomes of Sweet Ann (**Fig2C**, green triangles and squares, respectively) were more dissimilar from respective microbiomes of Manresa and Marquis (**Fig 2C**, blue and orange triangles and squares, respectively). The analysis of BC distances further indicates that both the rhizosphere and the roots bacterial microbiomes of Sweet Ann were significantly different from corresponding microbiomes of Manresa and Marquis (**Fig 2E**, Rhizo: R^2^=0.16, p-value=0.01, Roots: R^2^=0.09, p-value=0.001). The rhizosphere fungal microbiome of Sweet Ann was also significantly different from the fungal rhizosphere microbiome of Manresa and Marquis (**Fig 2D** and **2F**, Rhizo: R^2^=0.13, p-value=0.01). However, the root fungal microbiome of Sweet Ann was not significantly different from the root fungal microbiome of Manresa and Marquis (**Fig 2D** and **2F**). The analysis of the rhizosphere and the root microbiomes indicates that the highly susceptible strawberry cultivar, Sweet Ann, assembles a microbiome that is distinct from the lesser susceptible cultivars Manresa and Marquis.

**Fig. 2.**
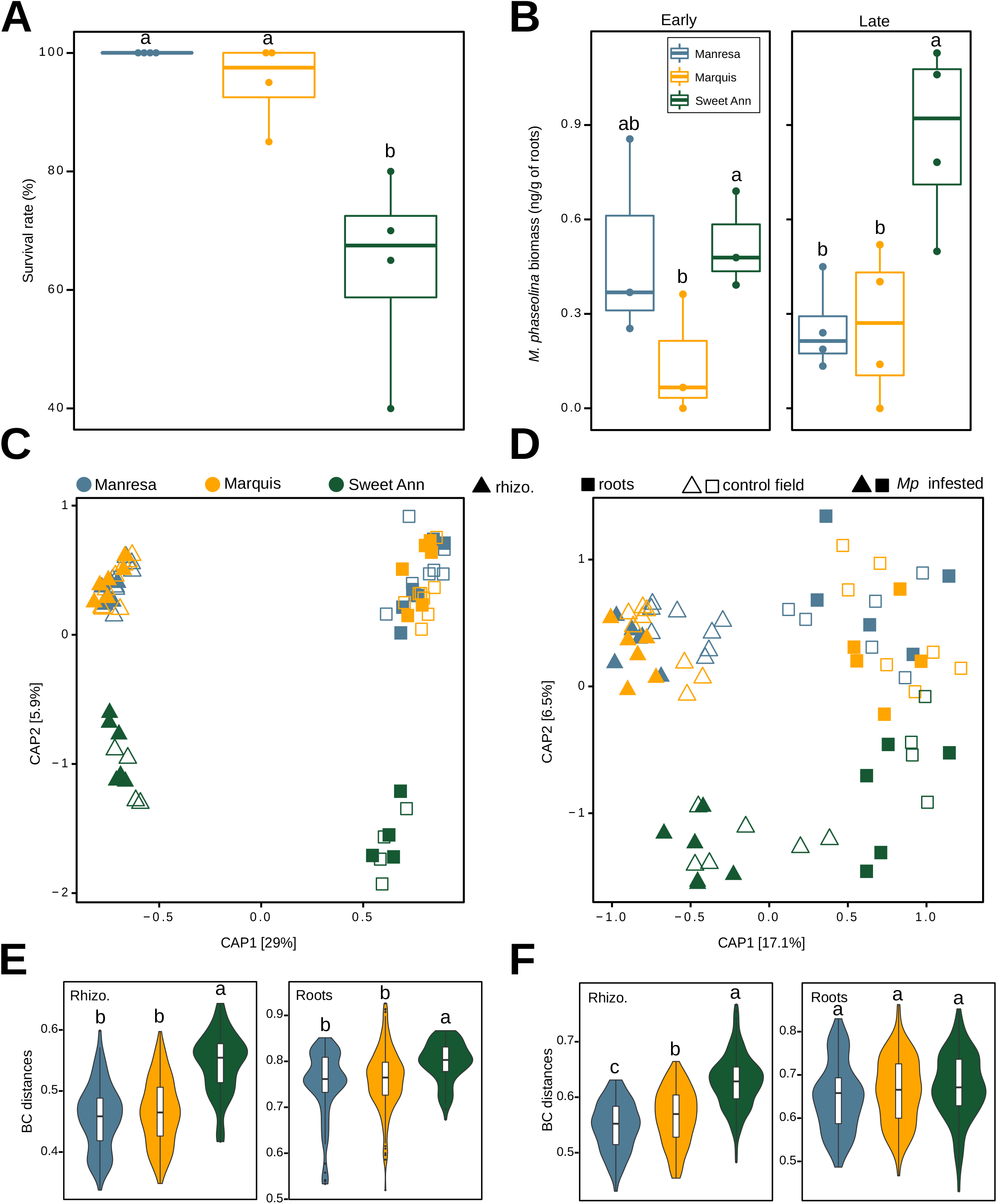
Highly susceptible cultivar, Sweet Ann, assembles a distinct rhizosphere and root microbiomes. **A**, the plot depicts survival rate (%) of strawberry plants planted in the infested field with *M. phaseolina* at harvest. The strawberry cultivar, Sweet Ann, shows lower survival rates to the fungal infection than Manresa and Marquis. **B**, box-plots show *M. phaseolina* biomass in nano-grams per gram of root tissue for either Manresa, Marquis or Sweet Ann at early (flowering) and late stage (harvest). **C and D** show constrained principal coordinate analysis of BC distances of bacterial and fungal communities associated with the rhizosphere and root samples of the three different strawberry cultivars, respectively. The structure of the root and rhizosphere bacterial microbiome Sweet Ann is more dissimilar from respective microbiome of Manresa and Marquis. Ffilled and unfilled shapes indicate *M. phaseolina* infested and control samples, respectively. **E** and **F**, violin plots displaying Bray-Curtis (BC) distances of bacterial and fungal communities in the three strawberry cultivars, respectively. Except of the root fungal microbiome, the structure of bacterial and fungal microbiomes of Sweet Ann are significantly different from those of Manresa and Marquis. Significance within-group medians were tested using Kruskal-Wallis significance and Conover’s multiple comparisons tests. Color-code is common over the plots and denotes strawberry cultivar.

To further reveal bacterial and fungal taxa that differentiate between the microbiome of the three strawberry cultivars, we fitted count data to zero-inflated Gaussian model and compared their relative abundance across the three cultivars (see **Methods**). Multiple bacterial and fungal taxa were preferentially enriched in the rhizosphere of Manresa and Marquis compared to Sweet Ann as depicted by ternary plots in **Fig 3A** and **3B**, respectively. The root bacterial microbiome of the three cultivars showed a similar trend but to a lesser extent (**Fig 3C**). Interestingly, 106 bacterial and 11 fungal ASVs were significantly enriched in the rhizosphere of both Manresa and Marquis, whereas Sweet Ann shared only one bacterial ASVs with Manresa and another bacterial ASV with Marquis (**Fig 3D** “upper right”). No fungal ASV was shared between Sweet Ann and any of the other test strawberry cultivars (**Fig 3D** “upper left). The root bacterial microbiome was overall less distinct and very few bacterial taxa were found to mediate differences between the three strawberry cultivars (**Fig 3D** “bottom left diagram”).

**Fig. 3.**
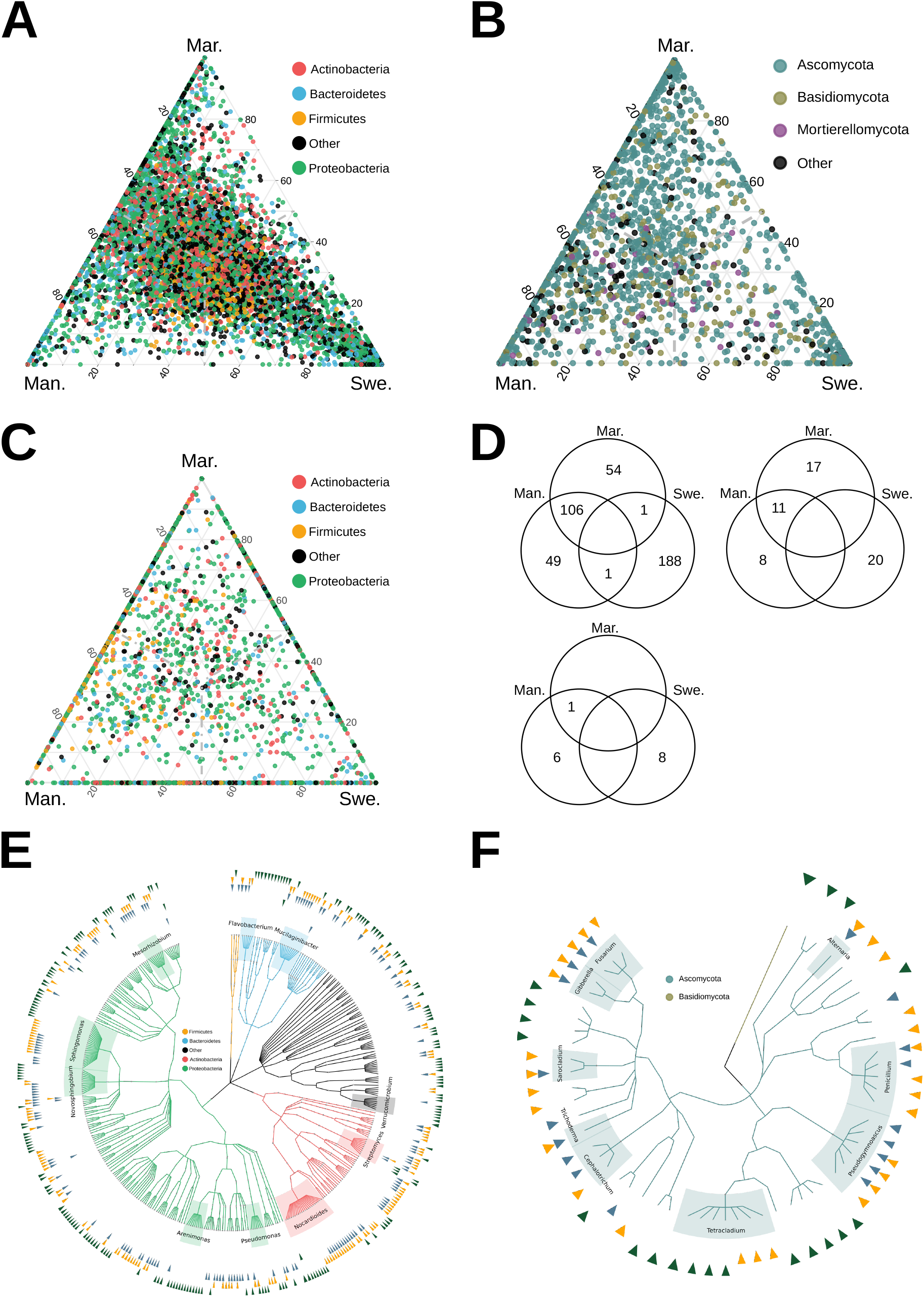
Differential enriched bacteria and fungi across the three strawberry cultivars. **A** and **B**, show ternary plots of the mean in relative abundance of rhizosphere bacterial and fungal ASVs, respectively. **C**, shows mean relative abundance of root bacterial ASVs. Circle indicates one ASV and color indicates the phylum. **D**, indicate venn diagrams of unique and conserved significantly enriched bacterial (top left) and fungal (top right) ASVs in the rhizosphere microbiome of the three cultivars, bottom left diagram shows unique and conserved significantly enriched bacterial ASVs in the root microbiomes. Digit in circles indicate number of enriched ASVs. **E** and **F** show taxonomic dendrograms of differently enriched bacterial and fungal ASVs, respectively. Color in tree branches indicates phylum. **E**, Arrow-head points to differentially enriched (D. E.) bacterial ASVs. Circle at 1^st^, 2^nd^ and 3^rd^ inner positions show D. E. ASVs in the rhizosphere of Manresa (blue), Marquis (orange) and Sweet Ann (green), respectively. Similar, circle at 4^th^, 5^th^ and 6^th^ positions indicate D. E. bacterial ASVs in the rhizosphere of respective strawberry cultivar. **F**, most inner, 2^nd^ and 3^rd^ circles indicate D. E. fungal ASVs associated to the rhizosphere of Manresa, Marquis and Sweet Ann, respectively. Prior to computing differential enriched ASVs, ASVs with less than 3 occurrences were filtered out, then count data were normalized using cumulative sum scaling and fitted to zero-inflated Gaussian mixture model. Significance is indicated by Benjamini-Hochberg adjusted p-value < 0.01. Color-code is common over the plots and denotes microbial phyla or strawberry cultivar as indicated in the graph.

By combining all results, more than half (220 ASVs) of the bacterial ASVs that mediated differences in the community structure among the strawberry cultivars belong to Proteobacteria (**Fig 3E, Supplementary-table 2**), whereas 98% of the fungal ASVs were Ascomycota (**Fig 3F, Supplementary-table 2**). Notably, several *Mesorhizobium, Sphingomonas, Novosphingomonas, Nocardioides*, and *Mucilaginibacter* ASVs were significantly depleted from the microbiome of Sweet Ann (**Fig 3E, Supplementary-table 2**). Comparably, the fungal microbiome of Sweet Ann was depleted from several ASVs belonging to *Fusarium, Gibberella, Sarocladium, Trichoderma, Cephalotrium, Pseudogymnoascus, Penicillium* and *Alternaria* (**Fig 3F, Supplementary-table 2**). Furthermore, Manresa and Marquis were commonly enriched with unique *Pseudomonas, Streptomyces* and *Tetracladium* ASVs that were depleted in Sweet Ann (**Fig 3E** and **3F**). All together, these results provide the evidence that highly susceptible Sweet Ann assemble a distinct microbiome and that Manresa and Marquis assemble more similar microbiomes.

### Lower connectivity in the co-occurrence network of Sweet Ann

We performed co-occurrence network analysis to explore potential co-associations between microbiome members, and reveal positive and negative associations between and among bacterial and fungal species (see **Methods**). Co-occurrence networks were computed using neighborhood selection model (Kurtz et al. 2015) by combining roots and rhizosphere microbiomes for each cultivar separately (**Supplementary Fig 3A, 3B** and **3C**, respectively). Network topology inspection revealed that the microbial co-occurrence network of Manresa (581 vertices, 4043 edges) and Marquis (582 vertices, 4280 edges) were more complex and connected than Sweet Ann (443 vertices, 2292 edges, **Supplementary table 3**). Although positive correlations were dominant in all three networks, the microbiome network of Sweet Ann had two times fewer negative correlations (5.7% vs 11.6% “Marquis” and 11.2% “Manresa”). To further reveal network topological properties, we computed several network parameters indicated in **Supplementary Table 3**. Higher clustering coefficients combined with lower average path length indicate more complex and more connected networks with potential strong interactions between microbes (Poudel et al. 2016; Guo et al. 2022). Here, we found that network modularity and average path length were higher in the microbial network of Sweet Ann compared to those of Manresa and Marquis and noted comparable average clustering coefficients between Manresa and Sweet Ann (**Supplementary table 3**). Our network topological analysis reveals that microbial co-occurrence networks of the strawberry cultivar Sweet Ann is less complex with potentially weaker ecological interactions between microbes.

To further reveal microbes that positively or negatively co-associate with *M. phaseolina*, we plotted co-occurrence sub-networks that depict all microbes that directly co-occur with *M. phaseolina* (“FASV66” in sub-networks, Fig **4A** “Manresa”, **4B** “Marquis”, and **4C** “Sweet Ann”). Remarkably, the microbial sub-network of Sweet Ann was less complex (5 vertices, 6 edges, **Fig 4C** and **4E**) than the microbial sub-network of Manresa (28 vertices and 63 edges, **Fig 4A** and **4E**) and Marquis (26 vertices and 53 edges, **Fig 4A** and **4E**). Examining the vertices properties of the three sub-networks, the fungal pathogen *M. phaseolina* (FASV66) co-associated with mainly fungal taxa in the microbial sub-network of Sweet Ann and these co-associated fungal taxa showed lower eigenvector centrality scores in the microbial full network of Sweet Ann (Supplementary **Fig 4F**). Whereas in the microbial sub-network of Manresa and Marquis, *M. phaseolina* (FASV66) co-associated with bacterial and fungal species that were well connected in their respective full microbiome networks (**Supplementary Fig 3D** “Manresa” and **3E** “Marquis”). Noteworthy, neither microbial species that co-associated with *M. phaseolina* (FASV66) in the sub-networks were low abundant nor rare taxa (**Supplementary Fig 3G, 3H, 3I**). These results prompt us to test whether *M. phaseolina* (FASV66) is more at the periphery of the microbial network of Sweet Ann compared to the two remaining strawberry cultivars. To this end, we computed within-module connectivity (Zi) and among-module connectivity (Pi) scores (Guimerà and Nunes Amaral 2005; Poudel et al. 2016). The analysis of topological features of microbial co-occurrence networks revealed that the microbiome network of Sweet Ann had less module hubs than Manresa and Marquis (**Fig 4D**). More importantly, the soil-borne pathogen *M. phaseolina* (FASV66) showed further low Pi and Zi scores in the microbial network of Sweet Ann compared to Manresa and Marquis that had higher Zi scores (**Fig 4D**). These results indicate that the pathogen *M. phaseolina* (FASV66) is further peripheral in the microbial network of Sweet Ann.

**Fig. 4.**
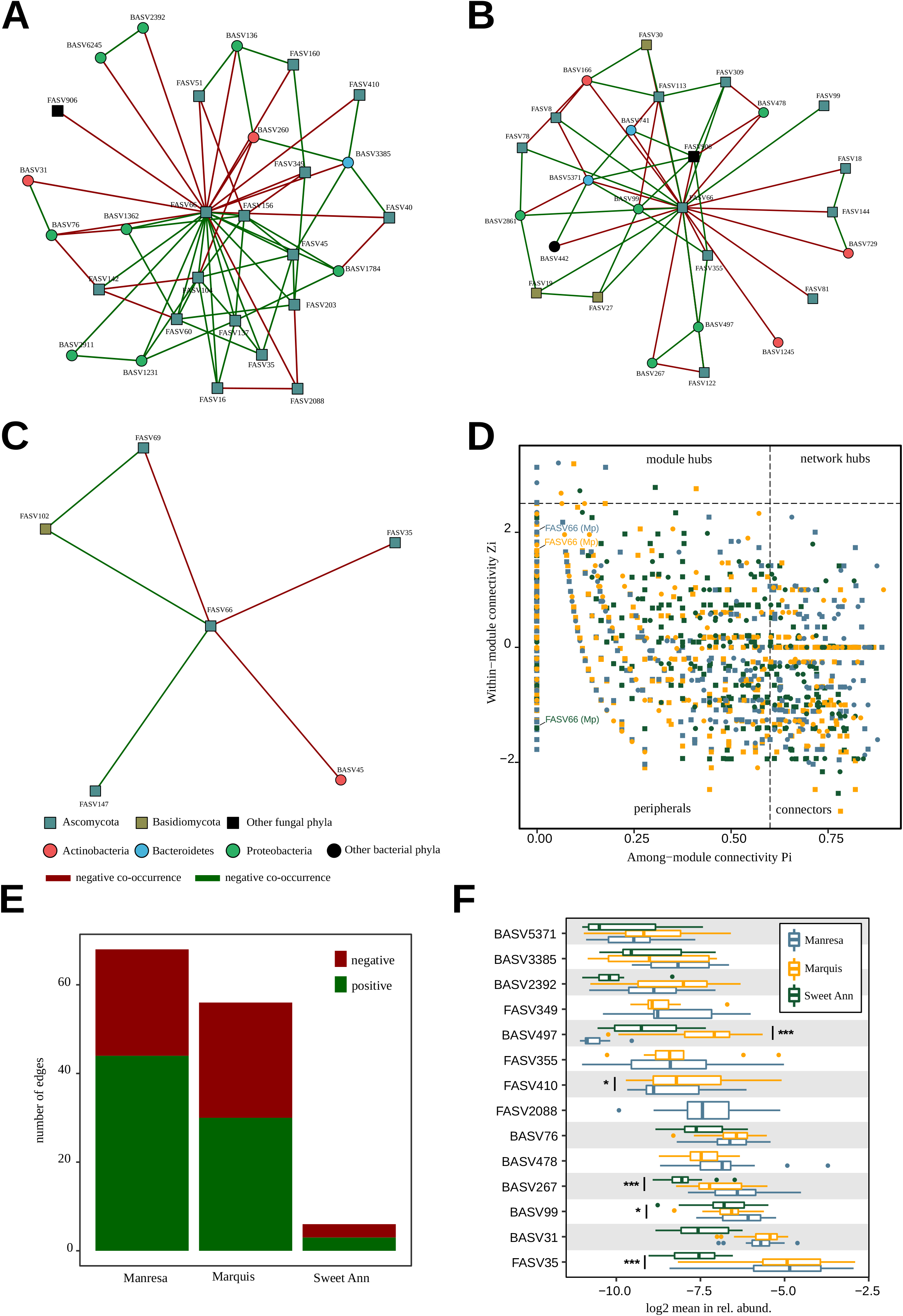
Connectivity in the microbiome network of strawberry cultivars. **A, B** and **C** show microbial co-occurrence sub-networks of bacterial and fungal species that positively or negatively co-associate with *M. phaseolina* (FASV66) in Manresa, Marquis and Sweet Ann, respectively. Co-occurrence networks were inferred using the neighborhood selection model by combining bacterial and fungal species associated to rhizosphere and root microbiome of each strawberry cultivar separately. Bacterial and fungi are indicated by circle and squarre vertices in the networks, respectively. Green and Red edges indicate postive and negative co-assocciation between two vertices and color in vertices indicates microbial phylum. **D.** Bi-plot shows within-module (Zi) and among-module (Pi) connectivity scores of bacterial and fungal species displayed in supplementary fig4. Vertices are divided into four categories based on Zi and Pi of 0.62 and 0.25, respectively (Guimera et al 2005). **E** shows total number of positive and negative edges in the sub-network of *M. phaseolina* (FASV66) indicated in panel A, B and C. **F** shows log_2_ mean in the relative abundance of selected bacterial and fungal species that negatively co-associate with *M. phaseolina* in all sub-networks displayed above. Color-code is common over the plots and denotes microbial phyla or strawberry cultivar as indicated in the graph. Significance within-group medians were tested using Kruskal-Wallis significance and Conover’s multiple comparisons tests.

Although these co-associations are inferred and might not always translate to ecological interactions (Blanchet et al. 2020), we wondered whether microbial species that negatively co-associate with *M. phaseolina* (FAS66) were comparably abundant in the microbiome of the three strawberry cultivars (**Fig 4F**). Comparing the relative abundance of these select bacterial and fungal taxa that negatively co-occur with *M. phaseolina*, four fungal (FASV349, *Albifimbria verrucaria*; FASV355, *Gibberella circinata*; FASV410, *Nectria ramulariae* and FASV2088, *Phoma schachtii*) and one bacterial (BASV478, *Luteibacter jiangsuensis*) ASVs were not recruited in the microbiome of Sweet Ann (**Fig 4F**). Although other bacterial taxa were recruited in the microbiome of Sweet Ann, they showed lower relative abundance in comparison to the microbiome of Manresa and Marquis (for instance BASV2392, *Sphingobium yanoikuyae*; BASV497, *Sphingobium aromaticiconvertens*; BASV267, *Devosia neptuniae*; BASV31, *Phycicoccus aerolatum* or BASV35, *Pseudarthrobacter* sp. - **Fig 4F**). Taken together, our network analysis shows that *M. phaseolina* is further peripheral in the microbiome network of Sweet Ann and showing fewer co-associates, that might indicate weaker ecological interactions within the microbiome of Sweet Ann.

## Discussion

The fungus *M. phaseolina* is a soil-borne pathogen that poses a significant threat to the strawberry cultivation. To reveal how the strawberry microbiome responds to *M. phaseolina* invasion and investigate links between disease resistance and microbiome composition, here we studied bacterial and fungal communities associated to the roots and the rhizosphere of strawberries upon infection. We compared the microbiome composition and structure of three strawberry cultivars with varying degrees of susceptibility (Sweet Ann, high; Marquis, moderate and Manresa, low) to *M. phaseolina.* We hypothesized that the rhizosphere and the root microbiomes of resistant cultivars assemble distinct bacterial and fungal communities that foster microbial connectivity. To this end, we profiled bacteria using 16S rRNA gene and fungi using ITS region in the bulk soil, the rhizosphere soil, and the roots of strawberry plants upon *M. phaseolina* infestation. Notably, our study shows that the root microbiome composition and structure of any tested strawberry cultivar was not significantly altered by the soil-borne pathogen (Bacteria; **Fig 1A**, Fungi; **Fig 1B, Table 1**). Despite observed shifts in the bulk soil microbial communities (**Supplementary Fig 1C, 1D**) and the rhizosphere fungal microbiome (**Table 1**), the root microbiomes were stable against *M. phaseolina* invasion (Bacteria; **Fig 1C**, Fungi; **Fig 1D**). To the best of our knowledge, this study is the first that investigates the response of the strawberry microbiome to *M. phaseolina* invasion. This finding is in contrast with several studies that have documented the response of phyllosphere microbiome to pathogen invasion (Agler et al. 2016; Zhang et al. 2019; Seybold et al. 2020; Durán et al. 2021; Li et al. 2022). For instance, Agler and colleagues studied the phyllosphere microbiome of thale cress, *Arabidopsis thaliana*, upon invasion by the oomycete pathogen *Albugo.* They showed that this oomycete pathogen significantly decreases microbial diversity and alters community structure in the phyllosphere of *A. thaliana* (Agler et al. 2016). Similarly, it has been shown that the wheat fungal pathogen, *Zymoseptoria tritici*, perturbs phyllosphere microbiome of the resistant wheat cultivar Chinese Spring (Seybold et al. 2020). Whether alteration of microbial communities upon pathogen invasion is a common feature is not clear. For instance, it has been shown that powdery mildew caused by *Erysiphe* negatively alters bacterial and fungal communities associated to the leaves of *Euonymus japonicus* (Zhang et al. 2019), whereas powdery mildew caused by *Golovinomyces orontii* alters only leaf fungi, but not bacteria, of *A. thaliana* (Durán et al. 2021). Our study showed that the root microbial communities of strawberries were unperturbed independently of their resistance phenotype and further suggest that strawberry plants assemble a root microbiome that is resistant to fungal perturbation. Future studies will help us understand whether this phenotype is shared across diverse strawberry cultivars and maintained under different abiotic conditions.

The rhizosphere and roots of plants are colonized by diverse microbial communities, these communities mainly derive from the adjacent soil (Bulgarelli et al. 2012; Lundberg et al. 2012) and extend the functional repertoire of the plant host (Bakker et al. 2018; Song et al. 2020). In this study, we performed a comparative analysis of the rhizosphere and root microbiomes of strawberry cultivars to better understand the interactions between the plant, the pathogen, and the microbiome. Our findings indicate that the highly susceptible strawberry cultivar, Sweet Ann, assembles a distinct rhizosphere and root microbiomes than the moderately or low susceptible cultivars (**Fig 2C, 2D, 2E** and **2F**). Enrichment analysis revealed that the differences between the microbiome of highly and moderate to low susceptible cultivars were explained by bacterial ASVs belonging mainly to Proteobacteria (*Mesorhizobium, Sphingomonas, Novosphingomonas, Arenimonas* and *Pseudomonas*), Actinobacteria (*Nocardioides, Streptomyces*) and Bacteroidetes (*Flavobacterium* and *Mucilaginibacter*, **Fig 3E**) and fungal ASVs belonging to Ascomycota (*Fusarium, Gibberella, Sarocladium, Trichoderma, Cephalotrichum, Pseudogymnoascus, Penicillium* and *Alternaia*, **Fig 3F**). These results rise the question whether the microbiome of Sweet Ann facilitated *M. phaseolina* infection (Stevens et al. 2021) or the microbiome of Manresa and Marquis mediated disease resistance. These two hypotheses are not mutually exclusive and could be tested in future experiments by performing reciprocal rhizosphere transplantation and evaluating disease incidence on strawberry plants (Kwak et al. 2018). It is well established that plants become enriched for beneficial microbes when challenged by fungal pathogens (reviewed in(Bakker et al. 2018; Li et al. 2021). In the frame of “Cry-for-Help” hypothesis, several studies have shown the potential of plants to recruit beneficial microbes from the untapped soil reservoir (Bakker et al. 2018). For instance, the root endosphere of sugar beet has been shown to recruit bacterial species belonging to *Chitinophagaceae* and *Flavobacteriaceae* during *Rhizoctonia solani* fungal infection. These beneficial bacteria were shown to harbor chitinase, nonribosomal peptide synthetases and polyketide synthetases gene clusters and suppressed the disease caused by the soil-borne pathogen *R. solani* (Carrión et al. 2019). Similarly, Chapelle and colleagues found the enrichment of several bacterial families (i.e. *Oxalobacteriaceae, Burkholderiaceae, Sphingobacteriaceae* and *Sphingomonadaceae*) in the rhizosphere microbiome of sugar beet when grown in disease-suppressive soil (Chapelle et al. 2016). Our finding suggest that Marquis and Manresa may recruit beneficial microbes that protect the strawberry plant from severe disease caused by *M. phaseolina*. However, above-mentioned studies employed disease-suppressive soils, the focus of this study was comparing the microbiome of strawberry cultivars with high or low susceptibility to *M. phaseolina*. Our observations join other studies that performed comparative microbiome analysis of resistant and susceptible plants to soil-borne pathogens (Kwak et al. 2018; Mendes et al. 2018; Lazcano et al. 2021). Collectively, these studies, including the present, suggest the importance of plant genetic traits in the assembly of the rhizosphere and root microbiomes that could mediate disease resistance and prevent pathogen invasion (Escudero-Martinez et al. 2022; Oyserman et al. 2022).

Prior to infecting the host plant, soil-borne pathogens encounter rhizosphere and root microbes. These microbial communities constitute first lines of defense against an invading pathogen (Chapelle et al. 2016; Kwak et al. 2018; Mendes et al. 2018; Carrión et al. 2019). Thus, understanding ecological interactions within microbiome members and between microbiome members and the plant pathogen is critical (Heijden and Hartmann 2016; Poudel et al. 2016). To gain further insights on potential ecological interactions between the fungal pathogen, *M. phaseolina*, and the rhizosphere and root microbiomes of strawberries, we constructed multi-kingdoms co-occurrence networks and measured positive and negative relationships between microbial species in Manresa, Marquis and Sweet Ann (see **Methods**). Analysis of network topological properties showed differences between the highly susceptible strawberry cultivar, Sweet Ann, and the low and moderate susceptible cultivars, Manresa and Marquis, respectively. Indeed, microbiome network of Sweet Ann was less complex and less connected (**Supplementary Table 3, Supplementary Fig 3C**) compared to the networks of Manresa and Marquis (**Supplementary Fig 3A** and **3B**, respectively). Lower connectivity was indicated by lower edges (2292 *vs* 4043 Manresa; 4280 Marquis), whereas graph density indicated lower complexity (**Supplementary table 3**). Further, the microbiome network of Sweet Ann showed fewer negative co-associations compared to Manresa and Marquis (5.7 *vs* 11.2 and 11.6 %, respectively, **supplementary Table1**). Of note, it has been shown that biotic and abiotic stresses can disrupt microbial networks (Gao et al. 2022; Li et al. 2022) and that negative co-occurrences are proposed to promote network stability (Coyte et al. 2015). Although microbiome networks from the three strawberry cultivars showed comparable clustering coefficients, average path length and modularity were higher in the microbiome network of Sweet Ann than in Manresa and Marquis (3.28 vs 3.16 and 3.12; 0.45 vs 0.39 and 0.4, respectively, **Supplementary Table1**). These topological properties reveal that the microbiome networks of Manresa and Marquis have higher level of complexity than the microbiome network of Sweet Ann. Previous studies have shown that highly connected and complex microbial networks could reduce the success of pathogen invasion (Mendes et al. 2018; Wei et al. 2018; Li et al. 2022). Our results are in line with previous findings and suggest protection against pathogen invasion could be mediated by increasing niche overlay, intensified antagonistic interactions or consumption of resources in the strawberry model (Poudel et al. 2016). Strikingly, the analysis of microbiome sub-networks indicated that *M. phaseolina* (FASV66) has very few connections in the sub-network of Sweet Ann (**Fig 4C**) compared to the sub-networks of Manresa and Marquis (**Fig 4A, Fig4B**, respectively). Analysis of within-module connectivity (Zi) and among-module connectivity (Pi) indicated that *M. phaseolina* is not hub species, but rather remains in the peripheral position of the microbiome network (**Fig 4D**). In contrast, an earlier study of the oomycete pathogen *Albugo* sp. has shown that this pathogen has central role in the phyllosphere microbiome network of *A. thaliana* and was validated as keystone species (Agler et al. 2016). This discrepancy could be explained by the nature of the pathogen, *Albugo* sp. being a foliar pathogen and *M. phaseolina* being soil-borne pathogen, and the microbial complexity of the rhizosphere microbiome compared to the leaf microbiome. Our network analysis revealed unique topological properties in the microbiome networks of strawberry plants with varying degree of resistance to the soil-borne pathogen, *M. phaseolina*. Furthermore, our study has shown that the cultivar Sweet Ann assembles a microbiome network that is less complex and less connected to the pathogen *M. phaseolina*. Whether microbial connectivity could mediate disease resistance in the test strawberry cultivars remain to be tested in future experiments

In conclusion, plants recruit microbial associates that constitute the first line of defense against an invading pathogen. Using the strawberry plant model, we showed that the soil-borne pathogen, *M. phaseolina*, does not perturb the root microbiomes of three different strawberry cultivars with varying degree of resistance to *M. phaseolina*. Microbiome comparative analysis indicated that the highly susceptible cultivar Sweet Ann assemble a distinct microbiome that shows poor network connectivity, whereas more resistant cultivar were enriched in potential beneficial microbes and showed higher network connectivity. Collectively, these results suggest of genetic traits in the plant host that could be involved in the assembly of beneficial microbiome members. Our study reinforces the eminent role of the plant microbiome as trait of selection in breeding programs and stresses further understandings of the genetic and biological mechanisms that mediate microbiome assembly. Uncovering these mechanisms will be key for future plant breeding programs and susceptibility in agriculture.

## Methods

### Plant material and experimental design

In this study, we carried out a field experiment with three strawberry cultivars exhibiting a gradient of resistance against the soil-borne pathogen *M. phaseolina* as determined in previous field trials (Lazcano et al. 2021). The trial was established on the California Polytechnic State University campus in San Luis Obispo, California (USA) between fall 2017 and summer 2018. The soil at both field sites was classified as Pachic Haploxeroll on a 0-2% slope with a clay loam texture, pH of 7.1, and 3% organic matter. The strawberry plants were grown in 1.6 × 2.5 m plots established in raised beds (0.3 m high). Plots contained 20 plants of the same cultivar and constituted the experimental unit. The plots were arranged in the field following a complete randomized block design with four blocks (i.e., four replicates) per cultivar.

### Soil infestation by *Macrophomina phaseolina*

*M. phaseolina* was not naturally present in the soil at the field site and had to be artificially introduced. Additional plots were established as a non-inoculated controls. Prior to inoculum introduction, the field was fumigated (392 kg ha^−1^ of a 50% chloropicrin + 50% methyl bromide solution) to ensure the removal of other soil-borne pathogens. Fumigation was carried out in 2015, two years before the start of this trial, since the effects usually last 2-3 years. Two weeks after transplanting, 5 g of *M. phaseolina* inoculum (20,500-25,200 CFU g^−1^) was applied to the soil-crown interface. *Macrophomina phaseolina* inoculum was produced from three field isolates collected from diseased plants in 2014 and 2015 following a procedure by Mihail (Mihail 1992). The density of viable microsclerotia within the cornmeal-sand inoculum was enumerated with direct plating of 10^−3^ to 10^−4^ serial dilutions onto NP-10 medium (Kabir et al. 2004) and ten replicate plates of each dilution were used. The plates were incubated in the dark at 30°C for 7 days before counting colony forming units. Bare-root strawberry transplants of the cultivars Marquis (intermediate resistance), Sweet Ann (low resistance) and Manresa (high resistance), were transplanted into the field trials in November 2017 and grown until 200 days after transplanting in June 2018, at the end of the typical growing season for this area of California. Fertilization was carried out through subsurface drip irrigation (fertigation) during the growing season and preplant slow-release fertilizer was banded in the bed 15 cm directly beneath plant lines for a total input of 112 kg N ha^−1^.

### Soil and plant samplings

Plant mortality was determined visually at flowering and harvest by assessing the percentage of dead plants in each plot. At these times we collected also a representative sample of 4 plants per cultivar and treatment (inoculated or control) to determine pathogen concentration in the roots and crown. Total genomic DNA was extracted using a DNeasy Plant Mini Kit (Qiagen, Hilden, Germany). Colonization of root and crown tissues by *M. phaseolina* were determined using quantitative polymerase chain reaction (qPCR) of the plant DNA extractions following Burkhard et al. (Burkhardt et al. 2018).

Sampling of plant roots, rhizosphere and bulk soil for microbiome analysis was carried out at flowering, and at the end of the harvest season. In each sampling, we randomly selected four plants within each plot avoiding dead or dying plants to reduce the confounding effects of the dead root tissue in the soil microbiome. Root samples were collected using hand trowels at a distance of 5 cm from the plant and to a depth of approximately 10 cm and then composited to obtain one sample per plot (Lazcano et al. 2021). A 750 cm^3^ bulk soil sample (without the presence of roots) from each block was also collected from the center of the strawberry bed (50 cm from the nearest plant) at a depth of 10 cm. Rhizosphere soil was collected by dry sieving roots on sterilized 500 μm sieves. Subsequently, a subsample of lateral roots wasm sieves. Subsequently, a subsample of lateral roots was collected using scissors and subsequently rinsed with PBS + 0.5% Tween-20 using an orbital shaker to remove the remaining soil attached to the roots. Roots were then rinsed in 1% NaOCl for 2 minutes to sterilize the root surface and subsequently rinsed with nanopure water. Soil and root samples were lyophilized and stored at −80 °C prior to DNA extraction for the study of prokaryotic and fungal diversity.

### DNA isolation and sequence data processing

DNA was extracted from a 250 mg subsample of soil or plant root tissue using the DNeasy PowerSoil Kit (QIAGEN, Venlo, Netherlands) following the manufacturer’s recommendations. DNA yield was measured using a NanoDrop 2000 (Thermo Fisher Scientific, Waltham, MA, USA). Samples with yield below 5 ng/μm sieves. Subsequently, a subsample of lateral roots wasL were extracted again prior to amplification. The V3-V4 region of the prokaryotic (including bacterial and archaeal) small-subunit (16S) rRNA gene was amplified with slightly modified versions of primers 338F (5’- ACTCCTACGGGAGGCAGCAG-3’) and 806R (5’- GGACTACHVGGGTWTCTAAT-3’)[29]. The 5’ ends of the primers were tagged with specific barcodes and sequencing universal primers. PCR amplification was performed in 25 μm sieves. Subsequently, a subsample of lateral roots wasL of reactions containing 25 ng of template DNA, 12.5 μm sieves. Subsequently, a subsample of lateral roots wasL of PCR premix, 2.5 μm sieves. Subsequently, a subsample of lateral roots wasL of each primer, and PCR-grade water. The PCR conditions for amplifying the prokaryotic 16S fragments comprised the following steps: initial denaturation at 98 °C for 30 seconds; 35 cycles of denaturation at 98 °C for 10 seconds, annealing at 54 °C/52 °C for 30 seconds, and extension at 72 °C for 45 seconds; and final extension at 72 °C for 10 minutes. PCR products were confirmed with electrophoresis in 2% agarose gel. Ultrapure water was used as the negative control to exclude false positives. PCR products were purified by AMPure XT beads (Beckman Coulter Genomics, Danvers, MA, USA) and quantified by Qubit (Invitrogen, USA). The size and quantity of the amplicon library were assessed with Agilent 2100 Bioanalyzer (Agilent, USA) and Library Quantification Kit for Illumina (Kapa Biosciences, Woburn, MA, USA), respectively. PhiX control library (v3) (Illumina) was combined with the amplicon library (at a fraction of 30%). The libraries were sequenced on Illumina MiSeq (2 × 300 bp) using the standard Illumina sequencing primers.

The ITS2 region of the eukaryotic (fungi) small-subunit rRNA gene was amplified with slightly modified versions of primers fITS7 (5’-GTGARTCATCGAATCTTTG-3’) and ITS4 (5’- TCCTCCGCTTATTGATATGC-3’) (Taylor et al. 2016). The 5’ ends of the primers were tagged with specific barcodes per sample and sequencing universal primers. PCR amplification was performed in a total volume of 25 μm sieves. Subsequently, a subsample of lateral roots wasL reaction mixture containing 25 ng of template DNA, 12.5 μm sieves. Subsequently, a subsample of lateral roots wasL PCR Premix, 2.5 μm sieves. Subsequently, a subsample of lateral roots wasL of each primer, and PCR-grade water to adjust the volume. The PCR conditions to amplify the eukaryotic ITS fragments consisted of an initial denaturation at 98 °C for 30 seconds; 35 cycles of denaturation at 98 °C for 10 seconds, annealing at 54 °C/52 °C for 30 seconds, and extension at 72 °C for 45 seconds; and then final extension at 72 °C for 10 minutes. The PCR products were confirmed with 2% agarose gel electrophoresis. Throughout the DNA extraction process, ultrapure water, instead of a sample solution, was used to exclude the possibility of false-positive PCR results as a negative control. The PCR products were purifyied using AMPure XT beads (Beckman Coulter Genomics, Danvers, MA, USA) and quantified by Qubit (Invitrogen, USA). The amplicon pools were prepared for sequencing and the size and quantity of the amplicon library were assessed using an Agilent 2100 Bioanalyzer (Agilent, USA) and with the Library Quantification Kit for Illumina (Kapa Biosciences, Woburn, MA, USA), respectively. PhiX Control library (v3) (Illumina) was combined with the amplicon library (expected at 30%). The libraries were sequenced either on PE300 MiSeq runs and one library was sequenced with both protocols using the standard Illumina sequencing primers, eliminating the need for a third (or fourth) index read. Raw bacterial and fungal sequencing reads have been deposited in Sequence Read Archive of the National Center for Biotechnology Information under BioProject number **PRJNA885096**.

### Sequences processing

Paired-end sequences obtained on the MiSeq platform were demultiplexed, followed by raw data filtering and processing utilizing the UPARSE pipeline (v11.0.667, www.drive5.com/usearch). Demultiplexed reads were screened to remove reads containing no call bases (N’s), screened and filtered to remove PhiX sequences and retain only reads with the proper primer pairs. Reads were then trimmed, overlapped, filtered to remove reads with overlapped length less than 300 bp, and then screened to remove reads containing Illumina adapter sequences using HTStream (https://github.com/s4hts/HTStream). Overlapped reads were denoised, summarized to Amplicon Sequence Variants (ASVs), and filtered for chimeric sequences using DADA2 (Callahan et al. 2016). Taxonomic assignment was done using the DADA2 implementation of the naive Bayesian classifier method, RDP (Wang et al. 2007a). Classification of 16S rRNA ASVs was done using the SILVA database reference (version 138.1) (Quast et al. 2012; Yilmaz et al. 2014). Classification of ITS ASVs was carried out using the UNITE database (version 04.02.2020) (UNITE Community 2019). ASVs were subsequently aligned using DECIPHER (version 2) and a phylogenetic tree was estimated using phangorn (version 2.5.5) in R (version 4.0.3) (Schliep 2011). Rarefaction curves were calculated for each sample using the vegan (version 2.5-6) package in R (version 4.0.3). Relative abundances of ASVs detected in at least four samples were included in subsequent analyses.

### Community composition and co-occurrence network analyses

Bacterial and fungal count tables, taxonomic tables and metadata tables used in this study are available as phyloseq objects at https://zenodo.org/record/7093889 . To compute alpha-diversity measures, Shannon and Observed ASVs (amplicons sequencing variants), count reads were rarefied to an even sequencing depth base of 1000 reads per sample for bacterial and fungal microbial communities using the R package “phyloseq” (McMurdie and Holmes 2013). Prior to compute Bray-Curtis (BC) distances, count reads were normalized by cumulative sum scaling normalization factors (Paulson et al. 2013). BC distances to centroid were computed using the multivariate homogeneity of groups dispersion available under the R function betadisper in the package “vegan” (v2.6-2, (Oksanen et al. 2022). To identify microbial ASVs that were differential enriched and test for their significance, ASVs count data were fitted to a zero-inflated Gaussian mixture model (Paulson et al. 2013) and ASVs showing Benjamini-Hochberg adjusted p-value < 0.01 were considered as significantly enriched. Constrained Principal coordinates analyses were computed using BC distances and constrained by the interaction terms, “cultivar” × “treatment” × “sample type”, using the R function capscale in the package “vegan” (v2.6-2, Oksanen et al 2022). Differences between group medians were tested using Kruskal-Wallis significance test and Conover’s multiple comparisons test, unless otherwise stated. All plotting graphs were displayed using the R package ggplot2 (v3.3.6, (Wickham et al. 2022)). Permutational multivariate analysis of variances in BC distances were computed using the R function adonis2 in the package “vegan” (v2.6-2, (Oksanen et al. 2022)). Relative abundance ternary plots were displayed using the R function ggtern in the package “ggtern” (v3.3.5, (Hamilton and Ferry 2018)). Taxonomic dendrograms were visualized using the GraPhlAn module (v1.1.4, (Asnicar et al. 2015)) Co-occurrence networks were inferred using the sparse inverse covariance estimation for ecological association inference pipeline and opting for neighorhood selection model as graphical model of inference (Kurtz et al. 2015). Networks were constructed by aggregated bacterial and fungal count data at the species taxonomic level and filtered out any species that had less than 6 occurrences across all samples for each strawberry cultivar separately using the R package “seqtime” (Faust et al. 2018). Networks visualization and calculation of network topological properties were done using R functions available in the package “igraph” (v1.3.1, (Csardi and Nepusz 2005)). Network modules were detected using the R function cluster_edge_betweenness in the package “igraph”. (v1.3.1, (Csardi and Nepusz 2005)). Within-module connectivity scores (Zi) and among-module connectivity scores (Pi) were computed as previously described (Guo et al. 2022) and the threshold value of 0.62 for Pi and 2.5 for Zi was followed as previously indicated in (Guimerà and Nunes Amaral 2005). All R scripts used in this study are available at https://github.com/hmamine/SRAM/tree/main/SRAM.

## Acknowledgments

We thank Dr. Quan Zeng and Dr. Blaire Steven for providing valuable comment on the early version of the manuscript. We are grateful to Eric Boyd who led the collection of rhizosphere and root samples from the field experiment, and Blythe P Durbin-Johnson who performed exploratory data analysis during the early stages of this manuscript. This work was supported by the California Agriculture Research Institute of the California State University (CSU ARI 19-03-008), and the California Strawberry Commission.

---

**Supplementary figure 1.**
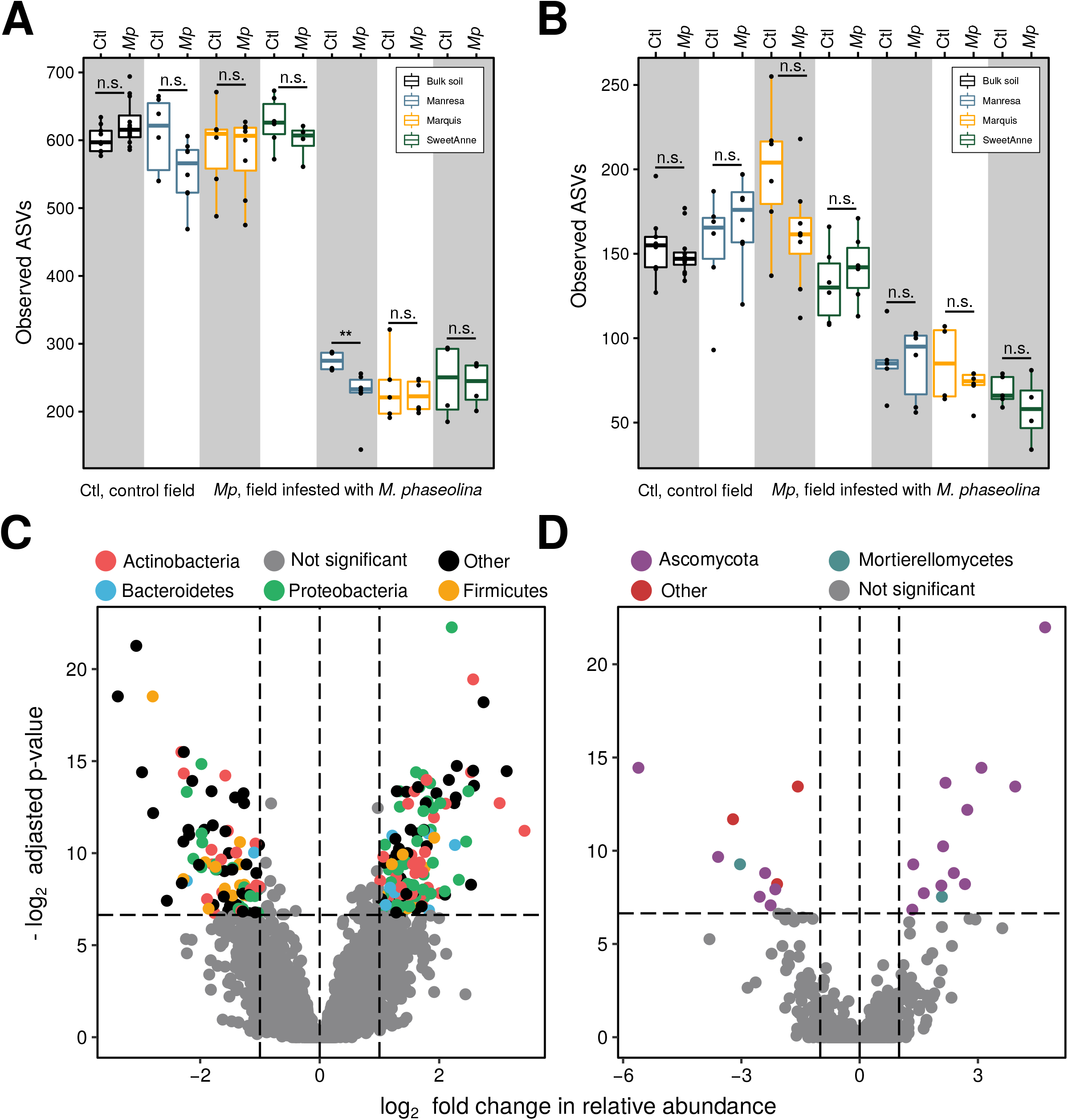
Panel **A and B** show box-plots that depict the alpha diversity measure, Observed ASVs of bacterial and fungal communities associated with different strawberry cultivars, respectively. Samples were rarefied to 1000 reads and significance difference was testing using Kruskal-Wallis test. Panel **C and D** show volcano plots that depict differentially enriched bacterial and fungal ASVs, respectively. Positive log_2_ FC indicates enriched in the control test condition, whereas negative log_2_ FC indicate ASVs that were enriched in the soil infested with *M. phaseolina*

**Supplementary figure 2.**
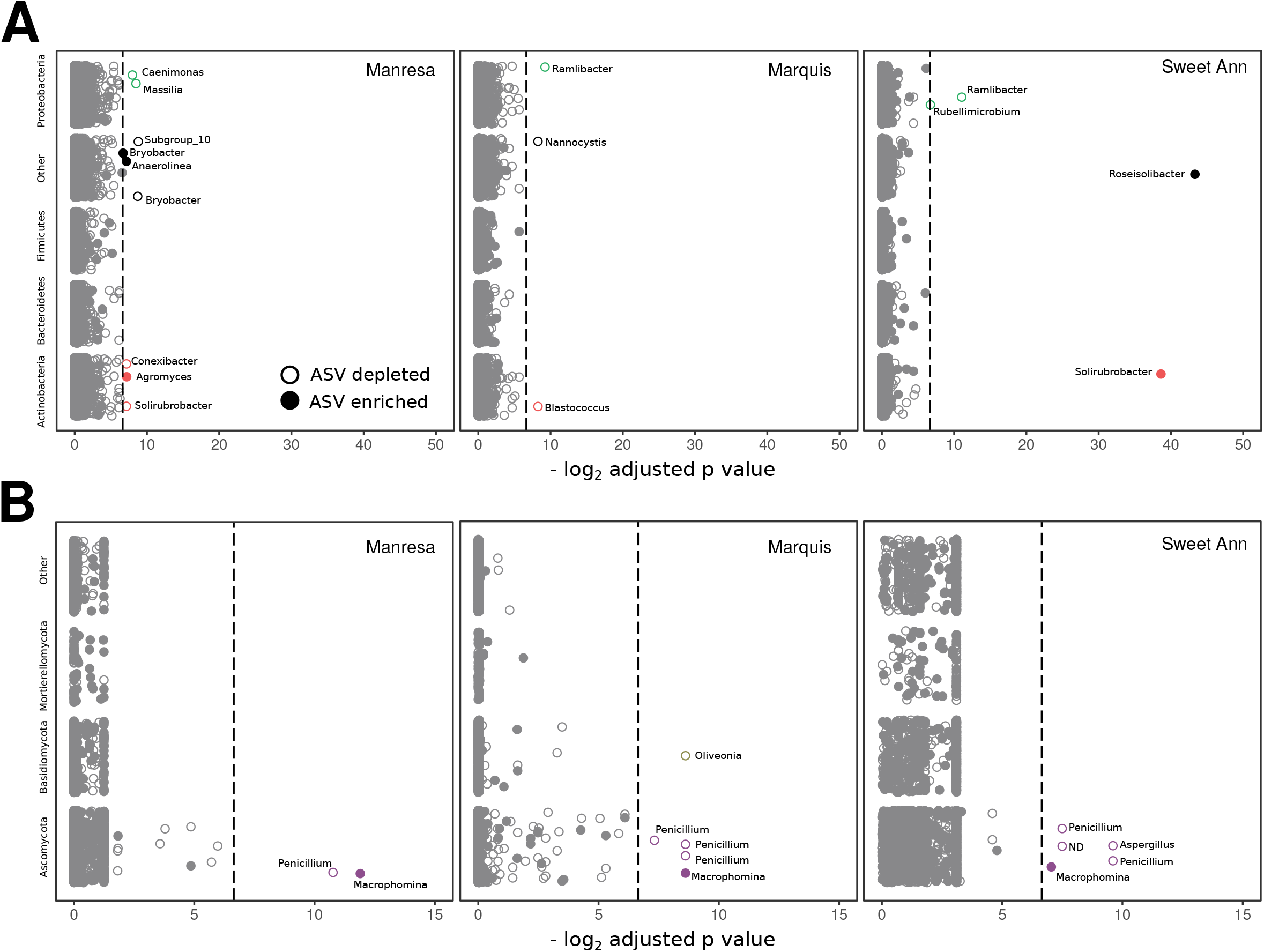
Panel **A and B** display Manhattan plot that depict −log2 Benjamini-Hochberh adjusted p-values of differentially enriched or depleted bacterial and fungal ASVs, in the rhizosphere of the three strawberry cultivars, respectively. Each circle indicates an ASV, color refers to the phylum and gray color shows not significantly enriched or depleted ASVs. Filled shape shows enriched ASV in the *M. phaseolina* condition and unfilled circle indicates depleted ASV from the control condition.

**Supplementary figure 3.**
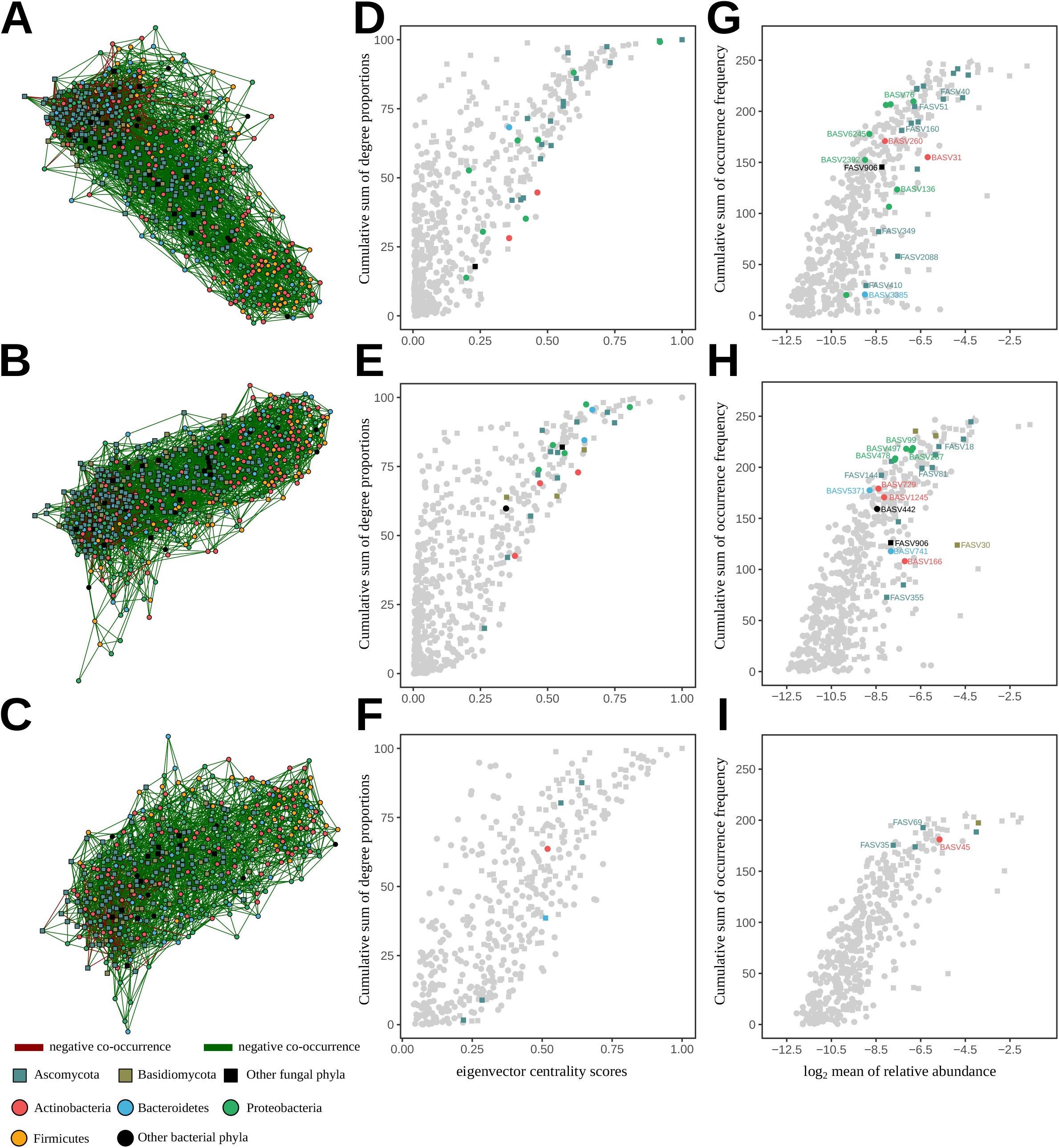
Panel **A, B** and **C** display bacterial and fungal co-occurrence network in the roots and rhizosphere samples of Manresa, Marquis and Sweet Ann, respectively. Bacterial and fungal ASVs were aggregated at Species taxonomic rank and species with less than 6 occurrences across all samples were filtered out. Each node represents a species and shape indicates kingdom. Species association networks were inferred using the neighorhood selection model by combining bacterial and fungal species associated to the roots and rhizosphere of each cultivar separately. Dark-green and dark-red edges indicate positive and negative co-occurrences. Scatter plots in **D, E** and **F** show node eigenvector centrality scores (hub score) function of the cumulative sum of degree proportions for each node in the network of Manresa, Marquis and Sweet Ann, respectively. Panel **G, H** and **I** indicate log_2_ mean of relative abundance of each node in the network function of its cumulative sum of occurrence frequency. Nodes that belong to *M. phaseolina* sub-network are highlighted according to their respective phylum color-code and nodes that negatively co-occur with *M. phaseolina* are further highlighted by their identification number.

## Notes

### Competing Interest Statement

The authors have declared no competing interest.

